# Nanopore sequencing to measure chromosome end-specific telomere lengths in human cells

**DOI:** 10.64898/2026.08.15.745035

**Authors:** Aljona Groot, Kayarash Karimian, Andreas Rechsteiner, Carol W. Greider

## Abstract

Telomere length has a significant impact on human health. Short telomeres cause age-related diseases, including pulmonary fibrosis, immunodeficiency, and bone marrow failure, while long telomeres predispose to cancer. Given the impact on human health, accurately measuring telomere length is important. A variety of methods have been developed over the past 40 years to measure telomere length. Many of these methods report only on the mean length of all of the telomere in the cell. Here, we describe the Telomere Profiling protocol using Oxford Nanopore Technologies (ONT) based on long read sequencing that can accurately measure chromosome specific telomere length.

## Introduction

Telomeres are specialized DNA–protein structures at the end of the chromosome, composed of simple GT rich repeats [1]. Telomeres shorten with each cell division, and the enzyme telomerase extends them to establish a distribution of lengths [2]. Short telomeres trigger DNA damage response and cellular senescence or apoptosis [3–5]. In humans, telomere shortening leads to stem cell failure age-related degenerative diseases, including pulmonary fibrosis, immunodeficiency, and bone marrow failure [6]. In contrast telomere elongation is essential for maintenance of cancer cell growth and inheritance of long telomeres predispose humans to cancer [7–9] The telomere length distribution was initially measured using Southern blots hybridized with telomere repeats to reveal the length distribution of all telomeres [10]. Other methods have been developed over the last 30 years to measure telomere length. Q-FISH uses fluorescence *in situ* hybridization (FISH) to estimate the telomere length on metaphase spreads [11, 12]. This method was able to capture the telomere shortening in telomerase deficient mice [13] but is only limited to measuring telomeres in metaphase spreads from actively dividing cells. A related method, Flow-FISH, measures the telomere content in white blood cells using flow cytometry [14]. This estimate of telomere length in a population of cells is highly reproducible and is the basis of the clinical assay to diagnose short telomere syndromes [14, 15], however it only measures length in human blood samples not other tissues. Quantitative PCR (qPCR) is a popular method used to assesses relative telomere content by comparing the amplification of telomeric DNA to a single-copy reference gene [16]. This method is easy to implement and has been widely used, however several groups have found very high variation between replicates which may make the assay less reliable [15, 17]. Another method, Single Telomere Length Analysis (STELA), is a PCR-based technique that offers insights into chromosome-specific telomere lengths at several specific telomeres [18]. There are also several short read whole genome sequencing based methods including TelSeq, Computel, qMotif, and Telomerecat which estimate total telomere content in a cell using short-read Illumina sequencing data [19–22]. While these short-read estimators can provide relative telomere length differences, they are sensitive to specific sequencing methodology and length estimation algorithms and thus report significant differences in telomere length estimates[23]. In addition, like Southern blots and Flow-Fish, these methods only report on a bulk mean length of all telomeres. Long read sequencing using PacBio was introduced to measure telomere lengths at specific chromosome ends [24], however the limited length of the fragments sequenced did not allow reliable mapping to all chromosome ends and limits the length of the telomeres that are measured [25]. To overcome the limitations of some of these methods, we developed Telomere Profiling using Oxford Nanopore Technologies (ONT) long read sequencing to determine telomere length and map reads to specific chromosome ends [26]. We initially developed nanopore sequencing to determine yeast telomere length using whole genome sequencing [27]. To adapt this method for measuring human telomeres, a physical enrichment step was added to maximize the number of telomeric reads [26]. Two other groups also measured telomere length in human cells using nanopore sequencing without physical enrichment [28, 29] and one group measured mouse telomeres [30] using a kit available from Oxford Nanopore Technologies.

## Results and Discussion

Telomere length is maintained around an equilibrium distribution; therefore, it is important to obtain many reads from each chromosome end to increase the statistical confidence in the telomere length determined for that end. To enrich for telomere reads we use the intrinsic telomeric 3’ overhang TTAGGG to ligate to a biotin-tagged adaptor (TeloTag) and purify the telomeres using streptavidin. The telomeres are released from the biotin tag by restriction enzyme digestion of the TeloTag, and after library preparation they are sequenced on an ONT MinION or Promethion flow cell.

We developed a bioinformatic workflow to measure chromosome end specific lengths. We filter for reads containing the TeloTag then use our algorithm, TeloNP, to determine the telomere-subtelomere boundary. TeloNP employs a rolling window to identify the discontinuity in the telomeric repeat patterns that demarcates the telomere-subtelomere boundary. The telomere length is calculated as the number of nucleotides between the subtelomere boundary and the TeloTag. The telomere reads are mapped against a custom reference genome that contains 500 kb of each chromosome end from which the telomere repeat regions have been removed to allow the unique subtelomere sequences more weight during alignment. After mapping and filtering for high-quality alignments, chromosome end-specific telomere lengths are determined for each chromosome end. We typically require at least 10 reads per end, however having more than 10 reads increases confidence in the telomere length.

There are three major differences between our method and those described by others. First, our method sequences both strands of the telomere, the G-rich strand and the C-rich strand, allowing us to account for sequencing biases in the two strands, and providing more reads. Second, our method physically enriches the telomeric DNA enabling sample multiplexing on a single flow cell. Third, we include the entire region containing telomeric repeats as part of the telomere.

When developing TeloNP to establish the telomere-subtelomere boundary, we specifically sought to include the chromosome proximal region, adjacent the subtelomere, that contains Telomere Variant Repeats (TVRs) [31, 32] in the telomere length calculation. These TVR sequences contain TTAGGG as well as variant repeats and so can be bound by TRF1 and TRF2 which are part of an equilibrium feedback mechanism involved in establishing telomere length [33]. Thus, from a biological perspective, it is important that the TVRs are measured as part of the telomere. Different groups have used different bioinformatic pipelines and different definitions of what sequences should be included as part of the telomere, leading to discrepancies in reported telomere lengths. A recently developed telomere length analysis algorithm, Telogator2 [34] employs a clustering approach using the TVR region to distinguish alleles.

Telogator2 measures both TVR length and the canonical TTAGGG repeat region length, yet in the final analysis, they report by default the 75^th^ percentile of the canonical repeat length as the telomere length. The TVR region can span up to 8kb on some chromosome ends, making the telomere length reported by canonical repeats significantly shorter at some chromosome ends when compared to algorithms such as TeloNP that includes the TVR (Figure 1A). In addition, given the relatively low read number and the often-wide variance at each chromosome end, makes using the 75^th^ percentile more sensitive to outliers than using the median. Thus Telogator2’s reported default telomere length is noisy and can be shorter or longer than those generated by TeloNP (Spearman correlation ρ = 0.726) (Figure 1A and 1B). In contrast, when Telogator2 TVR lengths and the canonical repeat lengths are added together and the median is used instead of the 75^th^ percentile, the results are highly concordant with telomere lengths determined by TeloNP (ρ = 0.993) (Figure 1C). A more recent algorithm, TARPON, also employs a windowing approach to count TVRs and canonical repeats to measure telomeres in long read sequencing data. TARPON gives the user the option to report the telomere length with the TVR region included or to report just the canonical repeats [35]. When TVRs are included for TARPON the telomere lengths are again highly concordant with TeloNP (ρ = 0.994) (Figure 1D). These strong correlations support the conclusion that different algorithmic approaches of TeloNP, Telogator2 and TARPON to detect the TVR-subtelomere boundary can yield concordant results when the TVR and canonical repeats are included. To avoid discrepancy in reported telomere lengths caused by different definitions of what sequences a telomere should include, as shown in Figure 1A, it is important to standardize the definition of the telomere-subtelomere boundary. The biological importance of TRF1 and TRF2 binding sites argues that the TVR region be included as part of the telomere, which will also allow simple comparison of lengths reported by different research groups.

**Figure 1.**
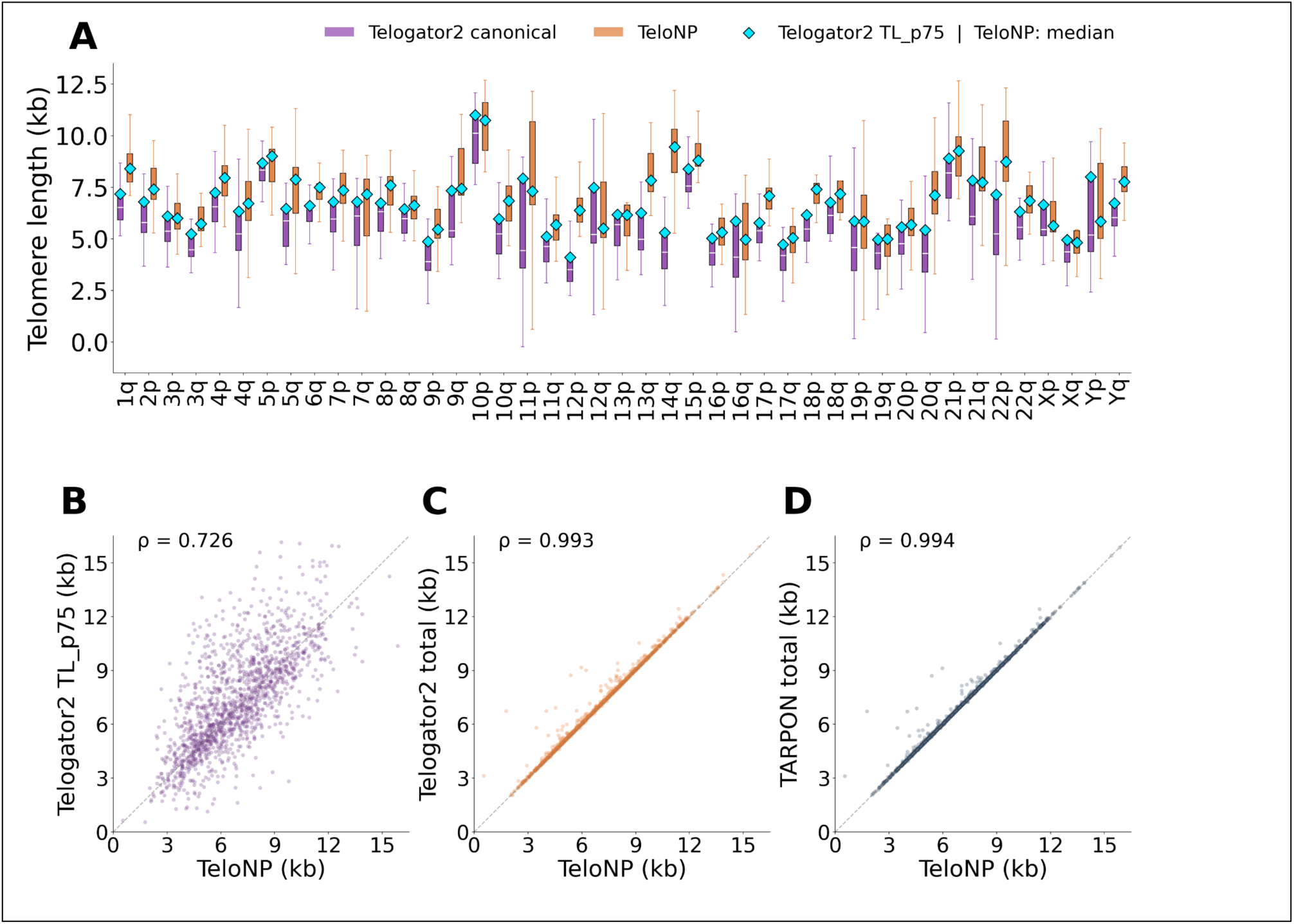
Telomere length estimates comparing TeloNP, Telogator2, and TARPON. Telogator2 requires ONT R10 flow cell data and so we used HPRC release 2 sequencing data for this analysis (<u>humanpangenome.org/hprc-</u> <u>data-release-2/</u>). Analysis was restricted to reads passing the TARPON multi-stage quality filter to ensure comparison across an identical read set for all tools. (**A**) Chromosome end-specific lengths are shown for Telogator2 and TeloBP. We used sample HG03521 and show 43 end that had two resolved alleles with ≥20 matched reads and only the higher-read-count allele was selected. Boxes show the read length distributions for Telogator2 canonical length only (purple) and TeloNP (orange) (median: white line; box: IQR; whiskers: 1.5×IQR). The Telogator2 reported canonical length, TL_p75 (75th percentile of the canonical lengths) and the TeloNP median lengths for each chromosome end are indicated by cyan diamonds. (**B–D**) Telomere length comparisons across 18 samples of the HPRC release 2 data set, requiring ≥10 reads per individual chromosome haplotype (n = 1,423 alleles). Dashed line: y = x. ρ: Spearman rank correlation. (**B**) the Telogator2 reported canonical length (TL_p75) vs. TeloNP (ρ = 0.726, median ratio = 1.01). (C) Telogator2 total length (TVR + canonical) median vs. TeloNP median (ρ = 0.993, median ratio = 1.00). (**D**) TARPON total length (TVR + canonical) vs. TeloNP median (ρ = 0.994, median ratio = 1.00).

## Materials and Methods

### Materials

#### TeloTag ligation and clean up

1. 100 ng/μl Telotag mix
2. 10X HiFi Taq DNA Ligase Reaction Buffer
3. Hifi Taq Ligase
4. DNA LoBind tubes
5. MicroAmp™ TriFlex Well PCR Reaction Plate
6. Veriti™ 96-Well Thermal Cycler
7. DNA LoBind tubes
8. HulaMixer™ Sample Mixer
9. DynaMag-2 magnet for 1.5 mL microtube
10. MyFuge 12 Mini Centrifuge
11. SPRIselect Beads (Beckman Coulter)
12. Ethanol solution: 80% ethanol/water (vol/vol)
13. 10X rCutsmart Buffer (NEB)
14. Qubit assay tubes
15. Qubit 3.0 fluorometer
16. Qubit double-stranded DNA (dsDNA) BR assay kit

#### Telomere Enrichment

1. Dynabeads: Dynabeads™ MyOne™ Streptavidin C1
2. HulaMixer™ Sample Mixer
3. DynaMag-2 magnet for 1.5 mL microtube
4. MyFuge 12 Mini Centrifuge
5. Binding Buffer: Dynabeads™ kilobaseBINDER™ Kit
6. Wash Buffer: Dynabeads™ kilobaseBINDER™ Kit
7. 10X rCutsmart Buffer
8. Elution buffer
9. Qubit assay tubes
10. Qubit 3.0 fluorometer
11. Qubit double-stranded DNA (dsDNA) HS assay kit

#### Library preparation and sequencing

1. NEBNext® Companion Module v2 kit: NEBNext FFPE DNA Repair Mix, Ultra II End Prep Enzyme Mix, NEBNext FFPE DNA Repair Buffer v2, Salt-T 4® DNA Ligase
2. Veriti™ 96-Well Thermal Cycler
3. 0.2mL 8-Strip PCR Tubes, Natural SnapStrip, Individual Dome Caps
4. DNA LoBind tubes
5. SQK-LSK114 Kit: AMPure XP Beads, Ligation Adapter, Ligation Buffer, Long Fragment Buffer, Elution Buffer, Sequencing Buffer, Library Solution, Flow Cell Tether, Flow Cell Flush
6. Protein LoBind tube
7. HulaMixer™ Sample Mixer
8. DynaMag-2 magnet for 1.5 mL microtube
9. MyFuge 12 Mini Centrifuge
10. 50 mg/mL UltraPure™ BSA
11. Qubit assay tubes
12. Qubit 3.0 fluorometer
13. Qubit double-stranded DNA (dsDNA) HS assay kit
14. R10.4.1 flow cell
15. MinION Mk1B

#### Software

1. MinKNOW (version v5.7.5) or later):https://community.nanoporetech.com/downloads
2. Dorado (version 0.3.1 or later):https://community.nanoporetech.com/downloads

## Custom code

1. Telomere-Read-Analysis: https://github.com/GreiderLab/Telomere-Read-Analysis
2. TeloBP/TeloNP: https://github.com/GreiderLab/TeloBP
3. ONT demultiplexer: https://github.com/GreiderLab/ont-demultiplex

### Methods

This protocol allows enrichment of telomeric fragments from high molecular weight genomic DNA (HMW gDNA) and sequencing to determine telomere length (Figure 2) HMW gDNA is first digested with a restriction enzyme to help reduce viscosity, then specialized barcoded biotinylated tags (TeloTags) are ligated onto the telomeres using multiple rounds of annealing and ligation in a thermocycler which increase the yield of ligated telomeres. The unincorporated adaptors are removed by size selection SPRI beads. The tagged telomeric DNA is then enriched using streptavidin-bound magnetic beads. The TeloTagged fragments are released from the streptavidin bead using a unique restriction enzyme site designed into the TeloTag. The telomere fragments are then processed using a Nanopore library prep with end fill-in and nanopore adaptor ligation and sequenced using a MinION Mk1B or Promethion P2 solo form Oxford Nanopore Technologies.

**Figure 2.**
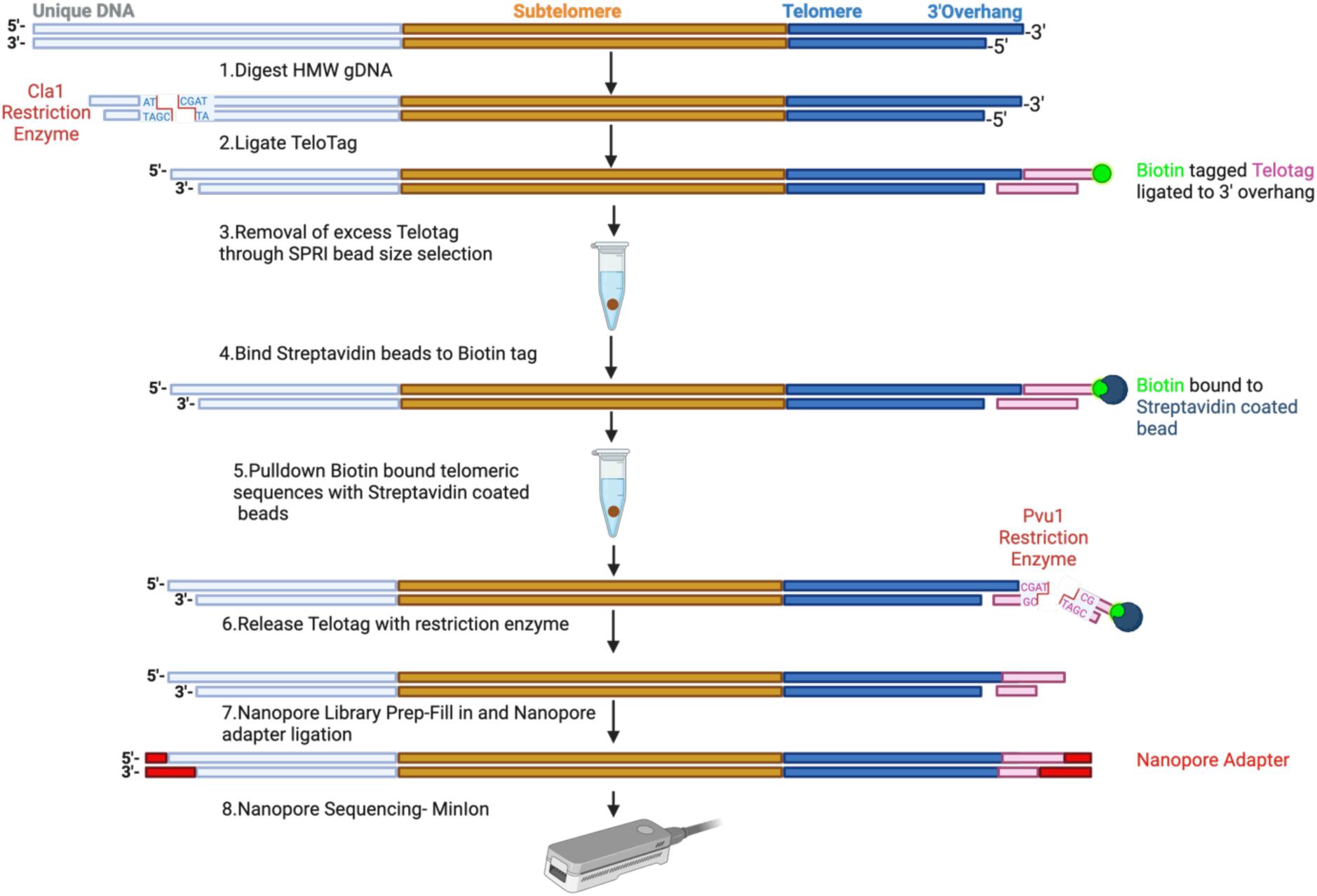
Overview of Telomere Profiling. HMW gDNA is digested with a restriction enzyme to reduce fragment size. The telomeric 3’ overhang is tagged with TeloTag using multiple ligation cycles, followed by removal of unligated TeloTag via SPRI bead size selection. The tagged HMW gDNA is then bound to streptavidin beads with gentle mixing, and the TeloTag is released from the beads by restriction digest. The enriched telomeres are processed through the Nanopore end-prep and sequenced using a MinION.

### Digestion of HMW gDNA for Telo Tagging

HMW gDNA is very viscous and if used directly will have a lower tagging efficiency. For this reason, the DNA sample is cut with a restriction enzyme to lower viscosity. Shearing is an alternative way to reduce viscosity; however, it was not explored in this protocol.

1. PureGene Gentra Kits maybe used to isolate HMW gDNA from whole blood, PBMCs, cultured cells, and tissues following the manufacturer’s protocol. This kit yields high-purity HMW DNA, typically 100–200 kb, A260/A280 ∼1.7–1.9. We also developed an alternate protocol is described in Note 10.
2. The optimal concentration of HMW gDNA is around 120–250 ng/µl. Prepare the digestion reaction as shown below in a DNA LoBind tube and mix gently with a wide-bore P200 pipette tip until the solution appears homogeneous.

| Content | Volume |
| --- | --- |
| HMW gDNA | 330 μl (40 μg) |
| 10X rCutsmart Buffer | 37 μl |
| ClaI | 3 μl (30units) |
| Total | 37 μl |

We recommend using 40 µg of HMW gDNA for robust chromosome end-specific telomere length measurement. For bulk telomere length analysis, the protocol can be adapted to use as little as 5-10 µg of HMW gDNA.

While other enzymes can be used, we found that cutting HMW gDNA using ClaI followed later by PvuI to release the telomeres, streptavidin beads provided robust results for chromosome end-specific telomere length measurement. When very little sample was available (< 10 ug), to increase telomere reads, we initially cut with BamHI and used EcoRI for release from the beads (see Section 3.4), as this combination yields shorter fragments and higher read counts. For analysis, we measured bulk telomere length (without chromosome-specific mapping).

1. Place the reaction mix tube in a heat block at 37°C for 2 hr. To ensure even mixing of the ongoing reaction, gently flick the tube every 30 min. Do not vortex. Place the tube on a heat block at 65°C for 20 min to heat inactivate the restriction enzyme.
2. The sample can be stored at 4°C for several months. Do not freeze.

### TeloTag Ligation

1. The telomere tagging reaction is prepared in 10 replicate 50 µl reactions using 4 μg of gDNA per reaction. This can be increased to 8 μg of DNA per PCR tube, if the concentration of HMW gDNA is higher than 120 ng/μl (*see* **Note 1**).
2. Using a 1.5 mL DNA LoBind tube, prepare the ligation mix for 10 tubes as described below

| **Content** | **Volume** |
| --- | --- |
| Digested HMW gDNA | 336 µl from previous reaction (40 µg) |
| HiFi Taq Ligase | 10 µl |
| 10X HiFi Taq DNA Ligase Reaction Buffer | 50 µl |
| TeloTag Mix (see Note 9) | 40 µl |
| Nuclease Free Water | 64 µl |
| Total | 500 µl |

Mix the sample gently using a P200 wide bore tip 20 times, or until the solution appears homogeneous. This step is critical. The sample will be viscous and mixing well ensures good distribution of components.

1. Aliquot the sample into ∼10 reactions of 50 μl each in a MicroAmp™ TriFlex Well PCR Reaction Plate. Seal the plate and briefly centrifuge at 200 rpm for 30 seconds to remove air bubbles.
2. Place the plate in a Veriti™ 96-Well Thermal Cycler and use the following cycling program.

First incubate the reactions at 65°C for 5 minutes. Perform 15 thermal cycles as follows:

i. Denature at 65°C for 1 minute.
ii. Anneal and ligate at 45°C for 3 minutes with a 15% ramp-down rate (∼0.78°C/min) between temperature transitions.

### Removal of unligated TeloTag by size selection

1. Place the bottle of SPRIselect beads on a HulaMixer™ Sample Mixer at 3 rpm for 1 h before setting up the reaction (*see* **Note 2**).
2. Combine the replicate ligation reactions into a 1.5 mL Low Bind tube. If multiple uniquely barcoded samples are being processed at the same time, they can also be combined at this step. Up to 6 uniquely barcoded reactions with 40 μg of gDNA may be combined. If the reaction volume exceeds tube capacity, gently mix reactions well before evenly splitting into separate tubes.
3. The pooled tagged DNA can be stored at 4°C for several days before the size selection step.
4. Aliquot the pooled sample into the appropriate number of reactions of 400 μl each in a 1.5 ml DNA LoBind tube. This is a volume sensitive step (*see* **Note 3**).
5. Add 0.45X (vol/vol) of the premixed and pre-calibrated (*see* **Note 2**) SPRI beads to your sample (180 μl beads for each 400 μl reaction). Use a new tip each time you dispense SPRI beads to assure correct volume is added (*see* **Note 2**). Do not pipet to mix the sample. Place the tube on the HulaMixer™ and invert mix the tubes with caps tightly closed for 20 minutes at 10 rpm.
6. Briefly centrifuge the tube to remove any beads from the cap of the tube. Place the tube on the DynaMag-2 magnet to isolate the beads. Wait for ∼2 min for the beads to concentrate at the magnet. Keeping the tube on the magnet, carefully remove the supernatant with a P200 pipette and save in a separate tube as it may be needed for troubleshooting. Rinse the beads with 1000 μl of freshly prepared 85% ethanol/water (vol/vol). Remove the ethanol by pipetting and then repeat the wash step. Remove the tube from the magnetic rack, centrifuge quickly, then place it back on the magnetic rack. Remove any residual ethanol by pipetting. Do not air dry the beads.
7. To release DNA from the SPRI beads, remove the tube from the magnetic rack and add 250 μl of 1X rCutsmart Buffer warmed to 65°C. Place the tube on heat block at 65°C for 20 min to release bound genomic DNA from the beads. Resuspend the beads by gently flicking the tube every 5 min. The heated elution step is essential for ensuring that long molecules are released from the beads. This can take up to 30 min.
8. Briefly centrifuge the tube to remove any beads from the cap of the tube. Place the tube on the magnetic rack, gently remove the supernatant with the bead free Telo tagged genomic DNA with a P200 and place it in a 1.5 mL DNA LoBind tube. If multiplexing, pool the reactions in the DNA LoBind tube. You can pool up to 600 μl of volume of size selected DNA in one 1.5 mL DNA LoBind tube.
9. Quantify the DNA concentration by using a Qubit BR Kit. The expected concentration is >100 ng/μl (*see* **Note 4**).
10. Quantify the DNA purity using nanodrop. The expected purity is A_260_/A_230_ > 2, A_260_/A_280_ > 1.8.
11. The tagged genomic DNA with the free TeloTags removed can be stored at 4°C for several months. Do not freeze.

### Telomere Pulldown and Enrichment

Telomere enrichment is a critical step in isolating telomere-specific DNA fragments from tagged and size-selected high molecular weight genomic DNA. The procedure involves binding DNA to streptavidin containing Dynabeads™, washing to remove any untagged DNA, and releasing the tagged telomeres from the biotin tag through restriction enzyme digest. Throughout the process, special care should be taken to avoid introducing air bubbles from pipet tips to maintain the integrity of the beads.

1. Place the Dynabeads™ MyOne™ Streptavidin C1 on HulaMixer™ Sample Mixer to mix at 3 rpm for 30 min before use.
2. Transfer 150 μl of resuspended Dynabeads beads per 40 ìg of starting gDNA in a Protein LoBind tube. If multiplexing, you can pool up to 600 μl of beads in one Protein LoBind tube, ensuring to not introduce air bubbles.
3. Place the tube on the DynaMag-2 magnet to isolate the beads. Wait for ∼2 min or until the solution has become clear. Carefully remove the supernatant with a P200 pipette.
4. Remove the tube from the DynaMag-2 magnet and add 125 μl (or if multiplexing half of the final volume) of Binding Buffer from Dynabeads™ kilobaseBINDER™ Kit.
5. Add the buffer directly on the beads. Use a P200 tip to resuspend the beads in the buffer and make sure not to introduce air bubbles.
6. Place the tube on the DynaMag-2 magnet to isolate the beads. Wait for ∼2 min or until the solution has become clear indicating that all the beads are held by the magnet. Carefully remove the supernatant with a P200 pipette.
7. Add 250 ìl of Binding Buffer from Dynabeads™ kilobaseBINDER™ Kit directly on the beads, or if multiplexing, add equal volume binding buffer as eluted DNA volume. Use a P200 tip to resuspend the beads in the buffer. If multiplexing, you may need to use more than one tube, make sure the maximum volume of beads gDNA and binding buffer does not exceed 1.4 mL for a single 1.5 mL Protein LoBind tube. Make sure not to introduce bubbles.
8. Add 250 μl of the TeloTagged gDNA to the resuspended beads using a wide bore P200 tip. Make sure not to introduce bubbles.
9. Place the tube on HulaMixer™ Sample Mixer to mix at 1 rpm at a 45° angle or taped to the so that the tube is “rolling” to gently mix for 20 minutes. Do NOT allow the tube to rotate end-over-end as this creates bubbles .
10. Briefly centrifuge the tube to bring the beads to the bottom of the tube. Place the tube on the DynaMag-2 magnet. Wait for ∼2 min or until the solution has become clear. Carefully remove the supernatant with a P200 pipette. Remove the tube from the magnet and add 1000 μl of Wash Buffer from Dynabeads™ kilobaseBINDER™ Kit. Use a P10 tip to displace the pellet from the side of the tube, and place the tube on HulaMixer™ Sample Mixer to mix at 1 rpm at a 45° angle for 5 min, without inversion
11. Repeat this wash step again.
12. Briefly centrifuge the tube to bring the beads to the bottom of the tube. Place on the DynaMag-2 magnet for ∼2 min until the solution clears. Carefully remove the supernatant with a P200 pipette. Remove from the magnet, add 1000 μl of Elution Buffer, and resuspend the beads on the HulaMixer™ at 1 rpm (45° angle) for 5 min. If multiplexing, sequentially elute beads from multiple tubes (up to 600 μl total) into one Protein LoBind tube using a wide-bore P1000. Repeat until all beads are combined into the same tube.
13. Briefly centrifuge the tube to bring the beads to the bottom of the tube. Put the tube on the DynaMag-2 magnet to isolate the beads. Wait for ∼2 min or until the solution has become clear. Carefully remove the supernatant with a P200 pipette. Remove the tube from the magnet and add 1000 μl of 1X rCutsmart Buffer. Place the tube on HulaMixer™ Sample Mixer to mix at 1 rpm at a 45° angle for 5 min for mixing. Do not invert the tube.
14. Briefly centrifuge the tube to bring the beads to the bottom of the tube. Place the tube on the DynaMag-2 magnet to isolate the beads. Wait for ∼2 min or until the solution has become clear. Carefully remove the supernatant with a P200 pipette. Remove the tube from the magnet and add 50 μl of 1X rCutsmart Buffer, and 3 μl (of the appropriate restriction enzyme to release telomeres from beads, see Note on restriction enzyme choice in **Section 3.1**).
15. Place the tube on heat block at 37°C for 30 min, and mix gently flicking the tube every 5 min. Do not allow the beads to settle at the bottom of the tube. Transfer the tube to a 65°C heat block for 20 min and mix gently flicking the tube every 5 min. Heated elution is essential for releasing long molecules from the beads. Never vortex tube.
16. Briefly centrifuge the tube and place on the DynaMag-2 magnet to isolate the beads. Wait for ∼2 min or until the solution has become clear. Carefully remove the supernatant with a P200 pipette, and place in a 1.5 mL DNA LoBind tube.
17. Quantify the DNA concentration by using a Qubit HS Kit. The expected recovery is approximately 0.1-0.01% of the starting gDNA sample (40-20 ng). For 40 μg of starting HMW gDNA, the expected recovery would be ∼10-15 ng of enriched telomeres (*see* **Note 5 and Note 6**).
18. The enriched telomeres can be stored at 4°C for several months.

## Library preparation and sequencing

This protocol is for library preparation and subsequent sequencing with an Oxford Nanopore MinION Mk1B and R10.4.1 flow cell. It is adapted from Oxford Nanopore Ligation sequencing DNA V14 (https://nanoporetech.com/document/genomic-dna-by-ligation-sqk-lsk114). Our method incorporates the barcodes as part of the TeloTag which is used to enrich the telomeres, and we have made small additional changes. If working with multiple libraries, multiplexing can be performed during the sample cleanup step using AMPure XP beads, with up to three libraries combined at this stage. Aim to load between 15 ng and 1 µg of your prepared library.

### Sequencing on Nanopore MinION Flow Cell

Follow the recommended Oxford Nanopore Ligation sequencing DNA V14 (https://nanoporetech.com/document/genomic-dna-by-ligation-sqk-lsk114) procedures for priming and loading the flow cell. You can expect ∼20-60% of the pores to be occupied at peak sequencing depending on the amount used. The N50 should be 35 – 50 kb. Depending on the sample purity and the original number of active pores in the flow cell, this step can take 1–2 d. You can use live basecalling or basecalling after the sequencing run. Check your computer requirements with the ONT guidance from MinION Mk1B IT requirements (https://nanoporetech.com/document/requirements/minion-it-reqs).

### Bioinformatic analysis of telomere length

We developed TeloNP to measure the telomere length of each sequencing read individually. TeloNP uses a rolling window to examine sequence content from the end of the telomere moving inwards and defines the telomere-subtelomere boundary as the transition from simple repeats to unique subtelomere sequence. The length of the telomere is calculated by TeloNP as the distance from the identified telomere-subtelomere boundary to the terminal TeloTag. TeloNP’s algorithm is robust against basecalling errors within telomere repeats common to Guppy and early Dorado basecallers. The TeloNP software is openly available at https://github.com/GreiderLab/. An alternative analysis package TeloGator2 [34] gives similar results when both the TVR and canonical repeat regions included in the length as described above. For high-accuracy datasets, such as those generated via the newest ONT chemistries and Dorado basecaller, TARPON is another effective method for telomere length determination if the entire telomere including TVR is reported [35]. The complete bioinformatic workflow for processing raw reads to determine chromosome-end-specific telomere lengths:

1. **Length filtering (optional):** To optimize downstream computational efficiency, an initial filter can be implemented to exclude reads shorter than 5 kb.
2. **Demultiplexing (optional):** Raw long reads can be demultiplexed by barcode into sample-specific FASTQ files utilizing a custom script available at https://github.com/GreiderLab. TARPON can also perform demultiplexing.
3. **Telomere length determination**: Read specific telomere lengths can be obtained by running TeloNP on the sample-specific FASTQ files. Alternatively, TARPON can be used to determine telomere length including the TVR regions.
4. Chromosome end-specific telomere lengths:

A custom reference genome is first constructed by isolating the terminal 500 kb from both ends of each chromosome within the chosen assembly. TeloNP is used to identify and remove the telomeres from the custom reference. Demultiplexed reads are mapped to this final telomere-depleted custom reference using minimap2. After filtering for high quality, primary alignments, chromosome-end-specific telomere lengths can be obtained, or the median length of all telomeres reported.

## Notes on Methods

### Note 1

Reaction setup for ligation of TeloTag to the telomeric end using. The starting HMW gDNA concentration should be in range of ∼120 – 250 ng/μl. The telomere tagging reaction can be split into multiple 50 µl reactions with 4–8 µg of HMW gDNA for each reaction.

When DNA is at a concentration of ∼120 ng/μl:

| **Content** | **Volume** |
| --- | --- |
| Digested HMW gDNA | 336 μl at 120 ng/μl |
| HiFi Taq Ligase | 10 μl |
| 10X HiFi Taq DNA Ligase Reaction Buffer | 50 μl |
| TeloTag Mix (see Note 9) | 40 μl |
| Nuclease Free Water | 64 μl |
| Total | 500 μl |

When DNA is at a concentration of ~250 ng/μl:

When DNA is at a concentration of ~250 ng/μl:
| **Content** | **Volume** |
| --- | --- |
| Digested HMW gDNA | 160 μl at 250 ng/μl |
| HiFi Taq Ligase | 10 μl |
| 10X HiFi Taq DNA Ligase Reaction Buffer | 25 μl |
| TeloTag Mix (see Note 9) | 40 μl |
| Nuclease Free Water | 15 μl |
| Total | 250 μl |

### Note 2

The bead suspension to sample ratio might vary with each preparation of beads due to evaporation from the SPRIselect bead bottle, or improper resuspension of the beads before use. Test the selection stringency of each bead preparation for removal of small DNA. A commercial DNA size ladder with a range of DNA fragment lengths should be tested to ensure the proper ratio of beads to DNA to the removal she correct sized small fragments. When dispensing SPRI beads, do not use the same tip more than once because of the high concentration of PEG, the volume dispensed will not be accurate after the first use.

### Note 3

This is a volume sensitive step. It is advisable to use a well calibrated pipettor, a P200 to ensure an accurate volume is being dispensed . Using a P1000 pipettor is not recommended. Instead, use a P200 to add 200 µl followed by 180 µl of beads for a total of 380 µl. A single error µl can result in sample failure.

### Note 4

DNA recovery can vary in this step. If the initial DNA concentration is lower than the recommended ∼114 ng/μl or there is an inaccurate measurement of the telomere tagged sample volume, it can reduce DNA recovery from the SPRIselect beads. HMW gDNA can also fail to elute from the beads during the 65°C incubation which may be avoided if the SPRIselect beads are tested for stringency of size selection (*see* **Note 2**). If the supernatants are saved in Section 3.3 TeloTag ligation, the SPRIselect bead cleanup can be repeated; to capture more bound genomic DNA from the beads and repeat the elution steps.

### Note 5

Low recovery can result from suboptimal telomere tagging, which often occurs when the HMW gDNA is too viscous during the tagging process, or from overly stringent wash steps that strip the tagged telomeres off the beads. You can stop and restart the HMW gDNA process, or you can proceed to library preparation if your recovery is within an acceptable range but consider pore occupancy will be greatly reduced when less DNA added.

### Note 6

Recovery that is unexpectedly high occurs because the wash buffer failed to remove the untagged DNA, which can happen if the HMW gDNA was too viscous. At this point, you can stop and restart the HMW gDNA process, or you can proceed to library preparation. Note that pore occupancy will be higher than expected.

### Note 7

If the solution remains viscous after overnight digestion, the hypotonic mix has not fully lysed the cells, and the proteinase K has not fully digested the proteins. Make 0.5X (5328 μl) of fresh lysis solution and hypotonic solution mix. Add the mix to the cells while vortexing at 6rpm.

Continue vortexing the sample every 15 min for 10 sec at 4-6 rpm until the solution is no longer viscous.

### Note 8

If the solution is not clear, you can do another round of protein precipitation by adding 0.17X (765 μl) of MPC Protein Precipitation Reagent (Lucigen), then repeat centrifugation. If you have already added more of the lysis and hypotonic solutions as described in Note 7, make sure the extra volume is accounted for in calculating the volumes of solutions needed for a second protein precipitation.

### Note 9

TeloTag design and annealing to generate double strand adaptor. The TeloTag consists of two single-stranded HPLC purified oligonucleotides, an adapter with a biotin tag on its 3’ end, a phosphorylated 5’ end, and a splint. The two oligonucleotides are annealed to generate the double stranded TeloTag which features a 3’ overhang to connect with the telomere, followed by an ONT barcode, restriction enzyme cut site, and biotin (Figure 3). We used ONT recommended barcode sequences (https://nanoporetech.com/document/chemistry-technical-document#barcode-sequences). We found empirically that specific barcodes worked better than others with the best ones being NB50, NB01, NB68, others that we used NB70 and NB72, NB69 gave good results. We did not test all of the barcodes and did not discover the reason for the variability. The splint is a set of 6 single-stranded oligonucleotides representing the six permutations of three telomere repeats of (CCCATT)_3_ that will anneal to any permutation of the telomeric 3’ overhang sequence. The barcoded TeloTag enables multiplexing of many samples. To generate the double-stranded TeloTag, two single-stranded oligonucleotides, the adapter, and the splint are annealed. The oligonucleotides are long to ensure complete sequencing of the captured telomere. The adapter contains a biotin tag on its 3’end, and the 5’end is phosphorylated to allow ligation with the telomere end.

**Figure 3.**
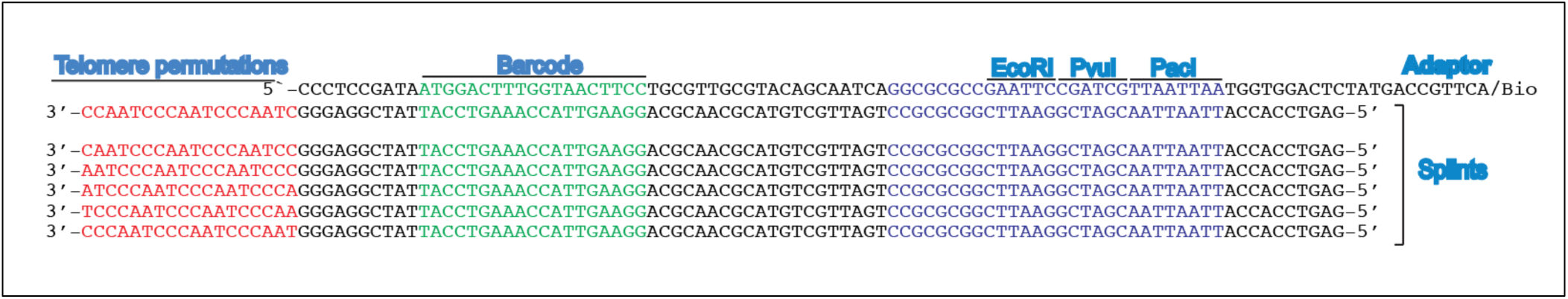
Sequence of a representative TeloTag adapter. The barcoded biotinylated adaptor (top strand) is annealed to a mixture of splints that have all 6 permutations of the CCCTAA telomere sequence to improve chances of in-frame annealing to the telomere 3’ overhang. The TeloTag includes a Nanopore-detectable barcode (shown in green), followed by three restriction enzyme cut sites (in purple). Additional sequence is included beyond the cut sites to ensure they are not at the very end of the molecule. The TeloTag was designed to be approximately 100 nucleotides long to minimize the loss of sequence information typically observed at the ends of nanopore reads.

Steps:

1. Resuspend the splints and adapter in nuclease-free water to a concentration of 100μM.
2. In each of a 0.2 mL 8-Strip PCR tube add 5 μl of each of the 6 splints for a total of 30 μl. Add 30 μl of the adapter.
3. Heat 10X HiFi Taq DNA Ligase Reaction Buffer to 65°C and vortex. Allow the tube to cool down to room temperature. Do not place on ice as salts in the reaction buffer will precipitate.
4. Add 10 μl of 10X HiFi Taq DNA Ligase Reaction Buffer, and bring the solution up to volume with nuclease-free water for a total of 100μl.
5. Mix the sample gently by pipetting. Use a thermocycler to anneal the splint and barcoded adapter to generate the double stranded TeloTag adaptor. The temperatures listed are for oligonucleotides that are ∼100 bp and have a Tm of (∼85°C). If different lengths are used, the time and temperature should be adjusted. The annealing step is done by heating the sample to 99°C and slowly decreasing the temperature 1°C /min until it reaches 85°C. Hold the temperature for 10 mins at 85°C, then cool to 4°C by decreasing the temperature 1°C /min until it reaches 4°C. This step is critical. If the tag does not anneal successfully, the telomeres will not be enriched.
6. The TeloTag stock can be stored at 4°C for several months.
7. Make a 1:100 working solution of the TeloTag by mixing 99 μl of water and 1μl of TeloTag, for a final concentration of 100 ng/μl. Aliquot and store at -20°C. Avoid freeze thawing the aliquots.

### Note 10

HMW gDNA Extraction without using commercial kit. The general extraction includes cell lysis, RNase treatment, protein precipitation, DNA precipitation (spooling). The DNA can be from cell lines, PBMC, or any other tissues. Starting with about 30 million cells a yield of 150 μg-300 μg of HMW gDNA should be expected. All reactions are kept on ice unless otherwise specified. Prepare all solutions using nuclease-free water (UltraPure DNase/RNase-free water).

## Steps

1. Collect 30 million cells from tissues culture cells or PBMCs through centrifugation in a 1.5 mL DNA LoBind Microcentrifuge tube by spinning at 300G for 10 minutes. Cell pellets can be flash frozen in liquid Nitrogen and stored at -80°C for later use.
2. Prepare fresh hypotonic solution on ice by mixing 990 μl of Nuclei Prep Buffer (Lucigen MasterPure Complete DNA/RNA Purification Kit) and add 33μl of 20mg/mL RNase A and 33 μl of 50U/μl RNase I_F_ (NEB)
3. Prepare fresh lysis solution on ice by mixing, 9 mL of the lysis solution from MasterPure Complete DNA and RNA Purification kit with 600 μl of 20mg/mL Proteinase K. If the solution becomes cloudy, the SDS has precipitated, warm the mix at 37°C until the solution is clear.
4. Thaw frozen cell pellet on ice.
5. Add 1056 μl of the hypotonic solution to the cells and resuspend the cell pellet by pipetting for 15-30 seconds.
6. Transfer the solution to a Falcon™ 50 mL High Clarity Conical Centrifuge Tube and keep on ice.
7. Add 9600 μl of the lysis solution, vortex to mix at 4 rpm for 10 sec (*see* **Note 7**).
8. Incubate the mixture in a 55 °C heat block for 4-6 hrs. Gently vortex the sample every 15 min for 10 sec at 4-6 rpm to ensure mixing.
9. Place the sample on ice for 5 min. If the solution turns cloudy, warm the mix at 37°C until the solution is clear. Return the sample to ice for an additional 5min.
10. Add 4.5 mL of MPC Protein Precipitation Reagent from the Lucigen/EpiCentre’s MasterPureTM Complete DNA and RNA Purification Kit A, while gently vortexing the sample at 4 rpm.
11. Centrifuge the sample at 2000 rpm for 30 min at 4°C. After 30 min the protein will have precipitated, and the supernatant will be clear (*see* **Note 8**).
12. Decant the supernatant into a DNA LoBind 50 mL Eppendorf® Conical Tubes. Precipitate the gDNA by adding 15 mL of cold isopropanol. Gently tilt the tube at a 45° angle ∼30 times, to allow for the DNA to precipitate and form a stringy cloud.
13. For clean HMW DNA spool the gDNA onto a glass pipette or remove it with a plastic pipette tip. Alternatively, gently pellet by centrifugation at 2000 rpm for 20 min at 4°C. Use a P1000 with the tip cut off to better transfer the DNA pellet to a new 1.5 mL DNA LoBind Microcentrifuge tube. Discard the supernatant.
14. Wash the pellet with 1000 ìl of freshly prepared 80% ethanol/water (vol/vol) and carefully remove the ethanol with a pipet tip. Repeat the wash step two more times. Pellet the DNA using a Mini Centrifuge. Remove any residual ethanol with a pipet tip. Air dry the pellet for 3 min.
15. Do not over dry the pellet, as this may affect DNA resuspension and recovery. This is a critical step. Over drying the pellet will make the DNA more difficult to resuspend.
16. Resuspend the pellet in 1000 μl of Elution buffer warmed to 37°C. Place the tube on a HulaMixer™ Sample Mixer at 37°C, to rotate at 1 rpm head over head overnight.
17. Quantify the DNA purity by using a nanodrop. The suggested purity is A_260_/A_230_ > 2, A_260_/A_280_ > 1.8.
18. Quantify the DNA concentration with a Qubit BR kit. The expected concentration is 100-300 ng/μl, which is a yield of 150 μg-300 μg total HMW DNA from 30 million cells. A low DNA concentration would be a result of insufficient cell lysing. See **Note 7**.
19. The HMW DNA can be stored at 4°C for several weeks. Do not freeze.

## References

1. Blackburn EH, Gall JG (1978) A tandemly repeated sequence at the termini of the extrachromosomal ribosomal RNA genes in Tetrahymena. J Mol Biol 120:33–53.

2. Greider CW, Blackburn EH (1985) Identification of a specific telomere terminal transferase activity in Tetrahymena extracts. Cell 43(2 Pt 1):405–413. 10.1016/0092-8674(85)90170-9

3. d’Adda di Fagagna F, Reaper PM, Clay-Farrace L, Fiegler H, Carr P, Von Zglinicki T, et al. (2003) A DNA damage checkpoint response in telomere-initiated senescence. Nature 426(6963):194–198. 10.1038/nature02118

4. IJpma A, Greider CW (2003) Short telomeres induce a DNA damage response in Saccharomyces cerevisiae. Mol Biol Cell 14(3):987–1001. 10.1091/mbc.02-04-0057

5. Lee HW, Blasco MA, Gottlieb GJ, Horner JW, Greider CW, DePinho RA (1998) Essential role of mouse telomerase in highly proliferative organs. Nature 392(6676):569– 574. 10.1038/33345

6. Armanios M (2022) The role of telomeres in human disease. Annu Rev Genomics Hum Genet 23:363–381. 10.1146/annurev-genom-010422-091101

7. de Lange T, Shiue L, Myers MR, Cox DR, Naylor SL, Killery AM, et al. (1990) Structure and variability of human chromosome ends. Mol Cell Biol 10:518–527.

8. Greider CW (1990) Telomeres, telomerase and senescence. Bioessays 12(8):363–369. 10.1002/bies.950120803

9. McNally EJ, Luncsford PJ, Armanios M (2019) Long telomeres and cancer risk: the price of cellular immortality. J Clin Invest 130:3474–3481. 10.1172/JCI120851

10. Larson DD, Spangler EA, Blackburn EH (1987) Dynamics of telomere length variation in Tetrahymena thermophila. Cell 50(3):477–483. 10.1016/0092-8674(87)90501-0

11. Rufer N, Dragowska W, Thornbury G, Roosnek E, Lansdorp PM (1998) Telomere length dynamics in human lymphocyte subpopulations measured by flow cytometry. Nat Biotechnol 16(8):743–747.

12. Zijlmans JM, Martens UM, Poon SS, Raap AK, Tanke HJ, Ward RK, et al. (1997) Telomeres in the mouse have large inter-chromosomal variations in the number of T2AG3 repeats. Proc Natl Acad Sci U S A 94(14):7423–7428. 10.1073/pnas.94.14.7423

13. Blasco MA, Lee H-W, Hande PM, Samper E, Lansdorp PM, DePinho RA, et al. (1997) Telomere shortening and tumor formation by mouse cells lacking telomerase RNA. Cell 91:25–34.

14. Baerlocher GM, Mak J, Tien T, Lansdorp PM (2002) Telomere length measurement by fluorescence in situ hybridization and flow cytometry: tips and pitfalls. Cytometry 47(2):89–99. 10.1002/cyto.10053

15. Alder JK, Hanumanthu VS, Strong MA, DeZern AE, Stanley SE, Takemoto CM, et al. (2018) Diagnostic utility of telomere length testing in a hospital-based setting. Proc Natl Acad Sci U S A 115(10):E2358–E2365. 10.1073/pnas.1720427115

16. Cawthon, R. M. (2002). Telomere measurement by quantitative PCR. Nucleic Acids Res, 30(10), e47. 10.1093/nar/30.10.e47

17. Aviv A, Hunt SC, Lin J, Cao X, Kimura M, Blackburn E (2011) Impartial comparative analysis of measurement of leukocyte telomere length/DNA content by Southern blots and qPCR. Nucleic Acids Res 39(20):e134. 10.1093/nar/gkr634

18. Baird DM, Rowson J, Wynford-Thomas D, Kipling D (2003) Extensive allelic variation and ultrashort telomeres in senescent human cells. Nat Genet 33(2):203–207. 10.1038/ng1084

19. Nersisyan L, Arakelyan A (2015) Computel: computation of mean telomere length from whole-genome next-generation sequencing data. PLoS One 10(4):e0125201. 10.1371/journal.pone.0125201

20. Lee M, Napier CE, Yang SF, Arthur JW, Reddel RR, Pickett HA (2017) Comparative analysis of whole genome sequencing-based telomere length measurement techniques. Methods 114:4–15. 10.1016/j.ymeth.2016.08.008

21. Farmery JHR, Smith ML, Lynch AG (2018) Telomerecat: a ploidy-agnostic method for estimating telomere length from whole genome sequencing data. Sci Rep 8(1):1300. 10.1038/s41598-017-14403-y

22. Holmes O, Nones K, Tang YH, Loffler KA, Lee M, Patch AM, et al. (2022) qmotif: determination of telomere content from whole-genome sequence data. Bioinform Adv 2(1):vbac005. 10.1093/bioadv/vbac005

23. Alder JK, Sutton RM, Iasella CJ, Nouraie M, Koshy R, Hannan SJ, et al. (2022) Lung transplantation for idiopathic pulmonary fibrosis enriches for individuals with telomere-mediated disease. J Heart Lung Transplant 41(5):654–663. 10.1016/j.healun.2021.11.008

24. Grigorev K, Foox J, Bezdan D, Butler D, Luxton JJ, Reed J, et al. (2021) Haplotype diversity and sequence heterogeneity of human telomeres. Genome research, 31(7), 1269–1279. 10.1101/gr.274639.120

25. Tham CY, Poon L, Yan T, Koh JYP, Ramlee MK, Teoh VSI, et al. (2023) High-throughput telomere length measurement at nucleotide resolution using the PacBio high fidelity sequencing platform. Nat Commun 14(1):281. 10.1038/s41467-023-35823-7

26. Karimian K, Groot A, Huso V, Kahidi R, Tan K-T, Sholes S, et al. (2024) Human telomere length is chromosome end-specific and conserved across individuals. Science 384(6695):533–539. 10.1126/science.ado0431

27. Sholes SL, Karimian K, Gershman A, Kelly TJ, Timp W, Greider CW (2022) Chromosome-specific telomere lengths and the minimal functional telomere revealed by nanopore sequencing. Genome Res 32(4):616–628. 10.1101/gr.275868.121

28. Schmidt TT, Tyer C, Rughani P, Haggblom C, Jones JR, Dai X, et al. (2024) High resolution long-read telomere sequencing reveals dynamic mechanisms in aging and cancer. Nat Commun 15(1):5149. 10.1038/s41467-024-48917-7

29. Sanchez SE, Gu Y, Wang Y, Golla A, Martin A, Shomali W, et al. (2024) Digital telomere measurement by long-read sequencing distinguishes healthy aging from disease. Nat Commun 15(1):5148. 10.1038/s41467-024-49007-4

30. Smoom R, May CL, Ortiz V, Tigue M, Kolev HM, Rowe M, et al. (2023) Telomouse—a mouse model with human-length telomeres generated by a single amino acid change in RTEL1. Nat Commun 14(1):6708. 10.1038/s41467-023-42534-6

31. Baird DM, Jeffreys AJ, Royle NJ (1995) Mechanisms underlying telomere repeat turnover, revealed by hypervariable variant repeat distribution patterns in the human Xp/Yp telomere. EMBO J 14(21):5433–5443. 10.1002/j.1460-2075.1995.tb00227.x

32. Coleman J, Baird DM, Royle NJ (1999) The plasticity of human telomeres demonstrated by a hypervariable telomere repeat array that is located on some copies of 16p and 16q. Hum Mol Genet 8(9):1637–1646. 10.1093/hmg/8.9.1637

33. Palm W, de Lange T (2008) How shelterin protects mammalian telomeres. Annu Rev Genet 42:301–334. 10.1146/annurev.genet.41.110306.130350

34. Stephens Z, Kocher JP (2024) Characterization of telomere variant repeats using long reads enables allele-specific telomere length estimation. BMC Bioinformatics 25(1):194. 10.1186/s12859-024-05807-5

35. Deimler N, Ho DV, Paul N, Gill Z, Baumann P (2026) TARPON—A telomere analysis and research pipeline optimized for nanopore. PLoS Comput Biol 22(2):e1013915. 10.1371/journal.pcbi.1013915

